# Evaluating the timecourses of morpho-orthographic, lexical, and grammatical processing following rapid parallel visual presentation: an EEG investigation in English

**DOI:** 10.1101/2024.04.10.588861

**Authors:** Donald Dunagan, Tyson Jordan, John T Hale, Liina Pylkkänen, Dustin A Chacón

## Abstract

Theories of language processing – and typical experimental methodologies – emphasize the word-by-word processing of sentences. This paradigm is good for approximating speech or careful text reading, but arguably, not for the common, cursory glances used while reading short sentences (e.g., cellphone notifications, social media posts). How can we interpret a sentence in a single glance? In an electroencephalography (EEG) study, brain responses to grammatical sentences (*the dogs chase a ball*) presented for 200ms diverged from non-lexical consonant strings (*thj rjxb zkhtb w lhct*) ∼160ms post-sentence onset and from scrambled constructions (*a dogs chase ball the*) ∼250ms post-sentence onset, demonstrating – at different time points – rapid recognition and cursory analysis of linguistic stimuli. In the grammatical sentences, unigram probability correlated with EEG data ∼150–300ms post-sentence onset, and probability of the word given its context estimated by BERT correlated with EEG data after ∼700–800ms. EEG responses did not diverge between grammatical sentences and their counterparts with ungrammatical agreement (*the dogs chases a ball*), although EEG responses did diverge for plural vs. singular morphology at ∼200ms. These results suggest that ‘at-a-glance’ reading is possible, based on coactivation of individual lexical items, morphological structures, and constituent structure at ∼200-300ms, but that words are not integrated into a coherent syntactic/semantic analysis, as evidenced by the substantially later responses to BERT probability and the absence of sensitivity to agreement errors.

## 1. Introduction

Most theories of language processing assume incremental word-by-word processing (e.g., Brasoveanu & Dotlačil, 2020; Frazier, 1987; Frazier & Fodor, 1978; Hale, 2001; Lewis & Vasishth, 2005). In the cognitive neuroscience of language, standard methodology follows this general framework – sentences are presented serially, word-by-word, with evoked brain responses measured at each word. This is motivated by the fact that speech, sign, and careful reading of longer texts are serial. However, seriality is a poor model for an increasingly common mode of reading: casual glances at short sentences (e.g., TikTok video captions, text notifications). The psycholinguistic mechanisms deployed in comprehending sentences read ‘at-a-glance’ are still poorly understood. With electroencephalography (EEG) and rapid parallel (∼200ms) visual presentation, we seek to: (1) establish when comprehenders distinguish grammatical sentences (*the dogs chase a ball*) from scrambled, ungrammatical counterparts (*a dogs chase ball the*) and non-linguistic consonant strings (*thj rjxb zkhtb w lhct*), i.e., isolate the ‘sentence superiority effect’ (SSE); and (2) establish the relative timecourse of this response to the interpretation of the lexical items in the grammatical sentences. We also sought to establish whether comprehenders build an integrated syntactic/semantic analysis of the sentence, in or subsequent to the SSE time period, by contrasting grammatical sentences with agreement violation counterparts (*the dogs chases a ball*). Our results suggest that the SSE depends on a mechanism that enables rapid detection of words and basic constituent structure. It is not the case that the analysis is detailed enough to reject agreement errors or to interpret words relative to their context until much later than the SSE.

Significant progress has been made in understanding the timecourse of word-by-word processing by associating event-related potentials (ERPs) with distinct stages of language processing (e.g., Friederici, 2011; Friederici & Kotz, 2003; Kaan, 2007). The early left anterior negativity (eLAN; generally observed over left anterior electrodes at 100–200ms) is associated with automatic local structure building, and can be found for word category violations (see Friederici & Weissenborn, 2007). The N170 and N250 (negative-going components observed over parietal and posterior sensors ∼200–300ms) are associated with initial visual processing and sub-word, morpho-orthographic processing (Grainger & Holcomb, 2009; Holcomb & Grainger, 2006, 2007; Morris & Stockall, 2012; Morris et al., 2008). A later left anterior negativity (LAN; at 300–500ms) is associated with case marking and morphosyntactic agreement (e.g., subject-verb number disagreement; see Molinaro et al., 2011). The N400 (a negative-going centro-parietal component ∼400ms) is associated with the access of lexical-semantic information (Friederici, 2011; Lau et al., 2008) and is highly correlated with surprisal (e.g., Frank et al., 2015; Lowder et al., 2018). Lastly, The P600 (a positive-going centro-parietal component between 500–900ms) is associated with the integration and retrieval of sentence-level syntactic and semantic information (Friederici, 2011). These ERPs – and their interpretations – importantly, are contextualized by word-by-word language comprehension; that is, the processing and integration of lexical items in a given parsing state. It is unclear to what degree these commonly-observed responses from serial reading and listening paradigms should be evoked in the reading of parallel input.

Several threads of research demonstrate that the visual system is capable of extracting information from more than a single word in one fixation. For example, skilled English readers are capable of detecting several letters to the right of a fixation (Rayner, 1998) and access some features of words in the parafovea (Antúnez et al., 2022; Rayner et al., 2003; Schotter et al., 2012). Therefore, it is possible that readers use parallel processing strategies, integrating information from across the visual field (Snell & Grainger, 2019). Similarly, findings from single-word processing studies also suggest rapid and parallel processing. A number of results suggest rapid, form-based processing of morphologically complex forms into their constituent parts (Rastle et al., 2000; see Taft & Forster, 1975). Magnetoencephalography (MEG) recordings show that morphologically complex forms (*re-made*; *sing-er*) exhibit increased activity in occipito-temporal regions ∼170ms (the ’M170’ event-related field; Zweig & Pylkkänen, 2009), and that this activity correlates with the relative probability of the whole word vs. the stem (Gwilliams & Marantz, 2018; Lewis et al., 2010; Solomyak & Marantz, 2011; see also Grainger & Holcomb, 2009; Holcomb & Grainger, 2006; Morris et al., 2013). This M170 response temporally precedes neural responses reflecting lexico-semantic interpretation of the word (Fruchter & Marantz, 2015; Pylkkänen & Marantz, 2003). This suggests that early stages of reading assess features of the form of multiple morphemes in a single fixation in single-word reading, which could in principle extend to short phrasal stimuli as well.

Nonetheless, it remains controversial how much linguistic information can be extracted from a single fixation, and how parallel these processes might be (Snell & Grainger, 2019; White et al., 2019). Recent work has explored the visual system’s capacity to extract grammatical and semantic information from short sentences read in a single fixation, presented rapidly, only visible for ∼200ms. Asano and Yokosawa (2011) found that Japanese readers were more accurate at recalling words in rapidly-presented sentences in which target words were contextually appropriate vs. sentences that had a semantic anomaly. To account for this, they propose that features of the sentence can be processed in parallel, and that a ’gist’ can be rapidly extracted. In a similar vein, in a series of studies in French sentences with 200ms presentation times, Snell, Grainger, and colleagues found an increase in word recall accuracy for grammatical and semantically plausible sentences vs. scrambled or implausible sentences (Massol et al., 2021; Snell & Grainger, 2017; Wen et al., 2019), which they refer to as the ’sentence superiority effect’ (SSE). In an (EEG) study, Wen et al. (2019) found that, compared to their scrambled counterparts, grammatical sentences presented in this fashion resulted in a reduction in amplitude of the N400, a component which is inversely correlated with the accessibility of lexical semantics in a context (Bornkessel-Schlesewksy & Schlesewsky, 2019; Kutas & Hilyard, 1984; Lau et al., 2008). Similar to Asano and Yokosawa (2011), Wen et al. (2019) propose that grammatical information is extracted in parallel from the fleeting stimulus, and that this information then interactively facilitates lexical access, resulting in a reduction in the N400 component.

With MEG, Dufau et al. (2024) – using the same stimuli and scrambled vs. grammatical comparison as Snell and Grainger (2017) and Wen et al. (2019) – localize a series of left-lateralized cortical regions that display an SSE. The first sensitive region is the inferior frontal gyrus (321–406ms), followed by the anterior temporal lobe (466–531ms), and then the inferior frontal (549–602ms) and posterior superior temporal gyri (553–622ms). Further MEG work elaborating on this paradigm reliably finds responses to grammatical manipulation even faster than 300ms. In English, Fallon and Pylkkänen (2024) found that grammatical sentences elicited greater activation than length-matched noun lists in left middle temporal cortex ∼125ms post-sentence onset. Similarly, Flower and Pylkkänen (2024) found sensitivity to grammatical sentences vs. reversed sentences starting at ∼210ms post-sentence onset, localized to a broad left fronto-temporal and temporo-parietal language network. These two MEG studies further demonstrate that the syntactic information gleaned from a single fixation may not be fully detailed. Fallon and Pylkkänen (2024) showed that the SSE is insensitive to agreement errors (*nurses cleans wounds*) and implausible thematic role reversals (*wounds clean nurses*), suggesting an analysis that is insensitive to word-to-word relations like subject-verb number agreement and thematic role likelihood. Their results demonstrate rapid detection of basic constituent structure ∼150ms, which we take to be distinct from the interactive lexical processes hypothesized by Wen et al. (2019). Similarly, Flower and Pylkkänen (2024) found that brain activity did not distinguish between grammatical sentences and sentences with two-word transpositions until ∼320ms, significantly after the earliest stages of neural SSEs. Finally, in an MEG study in Danish, Krogh and Pylkkänen (2024) replicated a neural SSE at ∼230ms, with activity localized to left inferior frontal and anterior temporal regions. Moreover, they observed that different features of well-formed grammatical sentences elicited neural effects at different times, with argument structure features impacting activity starting at ∼250ms, and yes/no questions involving a displacement diverging from declarative sentences at ∼500–720ms. These findings further support the claim that the SSE is dependent on early, top-down deployment of grammatical knowledge to identify basic constituent structure.

Here, we consider three distinct hypotheses about the timecourse of the SSE relative to other linguistic computations. We first seek to establish the SSE using EEG by contrasting Grammatical sentences (*the dogs chase a ball*) with their ungrammatical Scrambled counterparts (*a dogs chase ball the*). Having established the relative timecourse of the SSE, we then sought to determine where in the EEG topography and when in time this effect occurred relative to other ERP responses. We further contrast grammatical sentences to non-linguistic, character-matched Consonant Strings (*thj rjxb zkhtb w lhct*) trials and agreement violations (*the dogs chases a ball*).

Then, to query the relation between the SSE and lexical processes, we correlate the EEG responses recorded for the Grammatical trials with unigram probability ( *p*(‘dogs’) ) and probability of the word given its context ( *p*(‘dogs’ | ‘the chase a ball’) ). Contextual probability is estimated using BERT (Devlin et al., 2019), a bidirectional Transformer encoder (Vaswani et al., 2017) trained on the masked language modeling objective. Given one or more masked input tokens, the model learns to predict the corresponding vocabulary item(s), given the surrounding context. A trained BERT model assigns a probability for a word, given a bidirectional context. While BERT probability is context-dependent, unigram frequency is context-independent.

We consider the **morpho-orthographic processing** hypothesis, by which structure is identified by rapid visual detection of its visual correlates, such as short functional elements, like *the* aligned with the left edge of the visual field, or morphosyntactic markers such as nominal plural *-s* or verbal singular *-s*. This hypothesis predicts that the SSE should co-occur with earlier stages of orthographic processing, i.e., concurrent with the N170+N250 complex (Grainger & Holcomb, 2009; Holcomb & Grainger, 2006, 2007; Morris & Stockall, 2012; Morris et al., 2008), and analogizes the SSE with rapid ‘blind’ identification of morphosyntactic structure in vision (Rastle et al., 2008, Zweig & Pylkkänen, 2009; Solomyak & Marantz, 2010). To test this hypothesis, we contrast the grammatical sentences with non-linguistic, consonant string counterparts (*thj rjxb zkhtb w lhct*) to first identify the earliest stage of morpho-orthographic processing. The morpho-orthographic hypothesis predicts that the SSE should precede EEG responses correlated with unigram and contextual word probability as the mechanisms are proposed to operate on the visual form of the stimuli, prior to lexical access. Lastly, the morpho-orthographic hypothesis predicts that readers should be able to rapidly detect number agreement errors, since these are transparently reflected in the orthographic representation of English (i.e., *the dog**s** chase**s***).

Secondly, we consider the **constituent structure** hypothesis in which shortly after (or concurrent to) access of lexical items, comprehenders detect basic constituency, i.e., whether there is a subject noun phrase or verb phrase, without necessarily integrating the word meanings into a single interpretation, or checking formal relations such as agreement. We take this hypothesis to be compatible with Fallon & Pylkkänen’s (2024) findings of early detection of short sentences, regardless of the plausibility of their thematic role or agreement errors. This hypothesis predicts that the SSE is likely to occur after the N170+N250 complex, i.e. after the divergence between Grammatical and Consonant String conditions, and may be concurrent to or subsequent to activation of lexical items, i.e. unigram probability. However, this hypothesis also predicts that the SSE should be insensitive to agreement, i.e., there should be no point during the SSE in which the ungrammatical agreement and grammatical agreement trials diverge, especially if readers deploy top-down knowledge that enables ‘correction’ of minor errors (Flower & Pylkkänen 2024). Importantly, this hypothesis does not predict that there is insensitivity to morphology like the nominal plural marker -*s* or the verbal singular marker -*s*, but rather insensitivity to the ungrammaticality of illicit combinations. Finally, this hypothesis predicts that contextual integration of the lexical items should follow the SSE, i.e., the contextual probability correlations should occur substantially later than the SSE.

Finally, we consider the **coactivation hypothesis**, following Asano & Yosokawa (2011), Snell & Grainger (2017), and Wen et al. (2019). This hypothesis predicts that the SSE is coextensive with lexical access, i.e., grammatical structure facilitates lexical access in a parallel, interactive, cascaded fashion. Crucially, this predicts that the SSE should temporally co-occur with measures of lexical probability. Since this hypothesis links identification of the syntactic structure of the sentence with the lexical items, this hypothesis also proposes that effects of unigram probability should temporally co-occur or overlap with effects of conditional probability. It is unclear whether this hypothesis makes predictions about sensitivity to agreement, given that agreement is a formal relation that may not necessarily be activated by lexical processing. We also propose that this model predicts that lexical activation should be parallel, i.e., there should not be distinct ‘phases’ of processing each lexeme in the neural signal, but rather lexical probability of words should modulate the neural response at the same time.

## 2. Methods

### 2.1. Participants

Thirty-five self-identified English speakers were recruited from the ANONYMOUS CITY community. Thirty-three participants were right handed, as assessed by the Edinburgh Handedness inventory (Oldfield, 1971). All participants had normal or corrected-to-normal vision, and reported no history of language impairment. Participants gave written informed consent, and were compensated $10 per 30 minutes of participation time. The study was approved by ANONYMOUS UNIVERSITY (#ANONYMOUS NUMBER) and was performed in compliance with relevant laws and institutional guidelines.

### 2.2. Materials

We prepared 50 sets of sentences in four conditions: Grammatical, Scrambled, Ungrammatical, and Consonant String (Fig. 1A). All Grammatical trials (*the dogs chase a ball*) were simple transitive sentences consisting of five words: a determiner, an animate noun subject, a transitive verb, a different determiner, and an inanimate noun object. Half of the grammatical trials consisted of a plural subject noun ending in -*s* and a plural verb; the other half had a singular subject noun and a singular verb ending in -*s*. All words were monosyllabic and monomorphemic, with the exception of the plural morphemes -*s*. Scrambled sentences (*a dogs chase ball the*) were constructed by randomly permuting the words of the Grammatical trials, with manual verification that the new string did not form a new grammatical sentence. We did not exclude scrambled phrases that could create locally coherent sub-strings (e.g., the relative clause interpretation of *a ball the dogs chase*). Length-matched, unpronounceable consonant strings (*thj rjxb zkhtb w lhct*) were also generated. Ungrammatical trials were created from the Grammatical trials by either removing the required singular verb agreement morpheme -*s* from the verb (*the dog chase a ball*), or by adding an unlicensed singular verb agreement morpheme -*s* to the verb (*the dogs chases a ball*). Because number was counterbalanced across item sets, length was matched between the Grammatical and Ungrammatical conditions.

**Figure 1.**
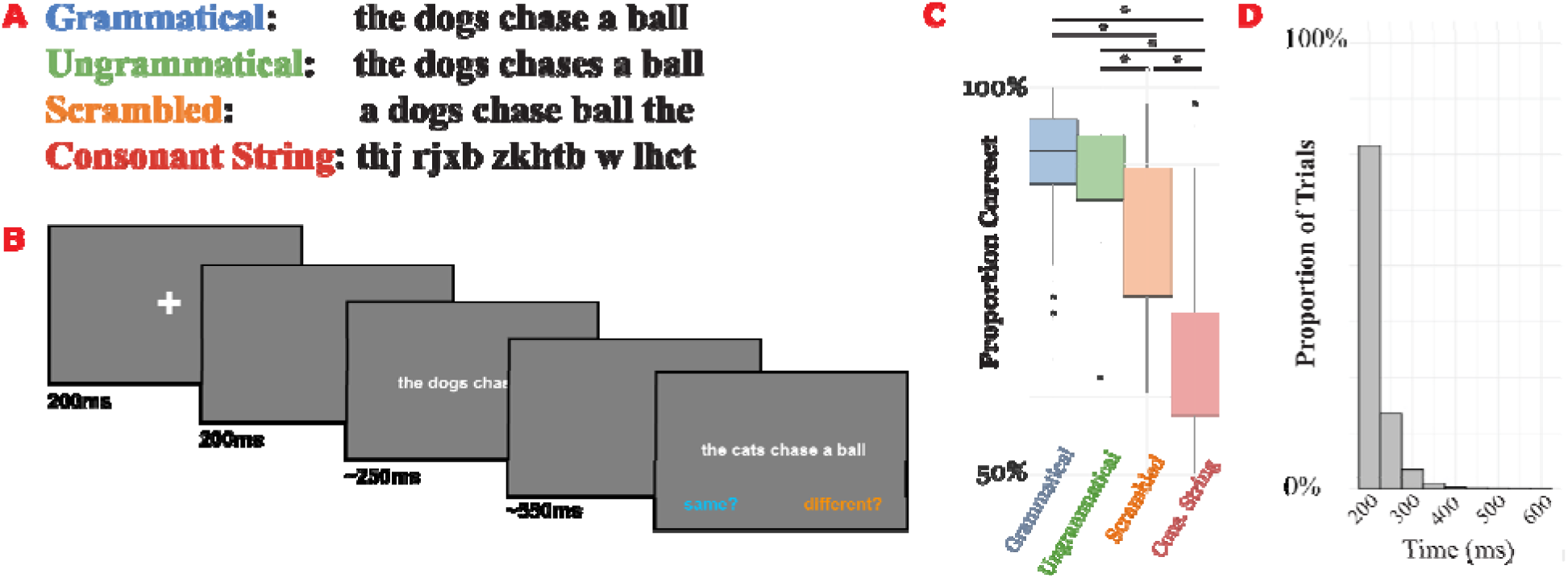
Experiment structure and behavioral results. (A) Example stimulus items for each condition. (B) Trial structure. (C) Accuracy by trial condition on the match/mismatch task. Stars signify a significant pairwise difference, as indicated by the logistic regression presented in §3.1. Each comparison is significant (*p* < 0.001), except for the comparison between Grammatical and Ungrammatical. (D) Proportion of trials by presentation time. 77% of trials were at the ceiling presentation time of 200ms.

Mismatch trials were constructed by replacing either the subject noun, the verb, or the object noun with a semantically similar length-matched foil (*the dogs chase a ball* ∼ *the cats chase a ball*). The probes in the match/mismatch task maintained all other features, i.e., scrambled word order, ungrammatical agreement, or consonant strings. In the Consonant String condition, the consonant string at either the subject noun position, the verb position, or the object noun position was replaced with another random consonant string of equal length.

Half of the stimuli were paired with a match trial, and the other half were paired with a mismatch trial. This was counterbalanced within-stimulus set, so participants could not leverage the identity of specific lexical items to guess whether a trial would be a match/mismatch. Instead, participants had to attend to the two nouns and verbs, i.e., the three content words.

### 2.3. Procedure

We used an adapted version of a match/mismatch task which has previously been employed in studies on behavioral and neural sentence superiority effects (Fallon & Pylkkänen, 2024; Flower & Pylkkänen, 2024, Krogh & Pylkkänen, 2024; Pegado & Grainger, 2020; Pegado et al., 2021). In this task, participants determine whether an untimed, second sentence is the same as, or different from, the rapidly displayed sentence.

All sentences were displayed centered on a screen, printed in a white monospace font against a dark gray background. Target sentences were displayed for a variable length, followed by a dark gray blank screen of variable length, which was adjusted such that the total time of the target sentence and the blank screen summed to 800ms. After the blank screen, participants saw another short sentence, which remained visible until participants pressed a button to indicate whether it matched. Participants entered their response using a keyboard, with the ’f’ key corresponding to ’match’ and the ’j’ key corresponding to ’mismatch.’ On-screen reminders (’same?’, ’different?’) were presented at each memory probe, to ensure participants remembered the mapping between key presses and responses. Participants were kept at a uniform distance from the screen, approximately 70cm from nasion to center of the screen. The visual angle subtended of the stimuli was approximately 15 degrees. The trial structure is shown in Fig. 1B.

Half of the trials were ’Mismatch’ trials, and the other half were ’Match.’ Unlike in prior RPVP studies using a match task (Fallon & Pylkkänen, 2024; Flower & Pylkkänen, 2024; Krogh & Pylkkänen, 2024; Pegado & Grainger, 2020; Pegado et al., 2021), sentence and blank screen display time were dynamically adjusted based on participant’s performance. This variable display time mechanic was implemented, in part, to conduct parallel experiments in other languages with different writing systems. We did not know *a priori* whether the minimum feasible stimulus length would be the same across writing systems and languages, and we expected by-participant variance, as well. Thus, we wanted the trial length to be adjusted dynamically to identify the ideal trial length for each subject. Sentence display time was constrained to vary between 200 and 600ms, initialized at 200ms. Sentence display time increased by 50ms after incorrect responses, and decreased by 50ms after correct responses. Participants received feedback on incorrect trials, with a 1000ms delay before the next trial. Faster presentation times were necessary to ensure that participants could not saccade during the critical time window, and thus read the stimuli serially. For our experiment, responses were remarkably accurate across trials (see § 3 and Fig. 1C). Thus, the average stimulus length was 216ms (SE = 0.43ms), and 77.0% of trials were at the minimum speed of 200ms (Fig. 1D), suggesting that variable display time should not be a substantial influence on performance or EEG responses.

Stimuli were presented in 4 separate blocks. Participants saw one trial per item set per block, with conditions and items evenly distributed between the 4 blocks. After each block, participants were instructed to take a break, which ranged from 2-5 minutes, and were provided with jokes to encourage them to not progress immediately into the next block. An experimenter was present with the participant during the recording and was available to answer questions or assist the participant if needed.

EEG signals were recorded using a 64 channel Ag/Cl BrainVision actiChamp+ system (Gilching, Germany). Impedance of the EEG sensors was reduced by the application of SuperVisc gel and lowered to <25kΩ. On-line EEG recording was referenced to FCz according to manufacturer standards, and then re-referenced to average sensors offline. Participants engaged in an unrelated task that is not reported here. The order of the two tasks was counterbalanced. Participants also engaged in a series of localizer tasks at the beginning of each recording session, but we do not report them here.

### 2.4. Data Availability

We make our data and analysis code publicly available via an Open Science Framework repository, the link to which has been removed for the sake of anonymity during the peer review process.

## 3. Results

### 3.1. Behavioral Results

Average accuracy was 81.42% (SE = 0.005). All participants except for one scored 70% or above. The data from all participants were analyzed. We used the lme4 package (v1.1-35.1, Bates et al., 2015) in R (v4.3.28, R Core Team, 2023) to fit a mixed effects logistic regression model to the accuracy data. The correct response was fit as the dependent variable, with condition as a 4-level factor, and with by-participant and by-item random intercepts. The model formula was Correct ∼ Condition + (1|Subject) + (1|Item). More complex random effect structures failed to converge. We fit the factor Condition with Grammatical as the intercept, since the research question at hand is how each sentence type diverges from the Grammatical condition. The data from all participants were analyzed. The results of the model are presented in Table 1. Afterwards, pairwise comparisons were conducted to assess which conditions diverged. Pairwise comparisons were conducted on the logistic regression model, with Tukey HSD correction, using the *emmeans* package (v1.10.5, Lenth, 2016). All conditions diverged from each other (*z*-ratios > 6, *p*s < 0.001), except for the pairwise comparison between Grammatical and Ungrammatical (β = 0.156, SE = 0.12, *z*-ratio = 1.35, *p* = 0.53), strongly suggesting that there is no detectable difference between sentences with grammatical agreement vs. sentences with agreement errors. The behavioral results are shown in Fig. 1C.

**Table 1.** Results of the mixed effects logistic regression fit to the behavioral responses. The model formula was Correct ∼ Condition + (1|Subject) + (1|Item). Bolded *p*-values are significant at alpha < 0.05.

| | $\beta$ | SE | $z$ | $p$ |
| --- | --- | --- | --- | --- |
| *(Intercept)* | 2.48 | 0.15 | 16.38 | **<0.001** |
| *Condition:*<br>*Ungrammatical* | -0.18 | 0.11 | -1.56 | 0.12 |
| *Condition:*<br>*Scrambled* | -0.83 | 0.10 | -7.93 | **<0.001** |
| *Condition:*<br>*Cons. String* | -1.73 | 0.10 | -17.37 | **<0.001** |

### 3.2. EEG Processing

All EEG preprocessing was conducted in MNE-Python (v1.4, Gramfort et al., 2013), and statistical analyses were conducted using Eelbrain (v0.39.5, Brodbeck et al., 2023). Raw EEG data was filtered off-line between 0.1–40Hz, using an IIR bandpass filter. We removed flat or noisy sensors (mean = 1.4, SE = 0.17 channels), and interpolated them using spherical spline interpolation (Perrin et al., 1989).

EEG data were re-referenced to an average reference off-line. Afterwards, epochs were extracted from –100ms to 800ms post-stimulus onset, i.e., the time of the rapid sentence presentation and the subsequent blank screen. The 100ms pre-onset period was used for baseline correction. Independent component analysis (ICA) was used to identify semi-regular endogenous electromagnetic noise sources, including eyeblinks, eye movements, and heartbeats. These components were removed. Following ICA, we automatically rejected all epochs that exceeded a 100μV peak-to-peak threshold. Afterwards, we visually inspected and removed other problematic epochs.

For each participant, we equalize the number of trials for each condition. Thus, each participant ends up with an equal number of good epochs for each trial condition. We then created condition averages for each participant for factorial analyses. After exclusion of the 18.4% of incorrect trials, exclusion of the unusable epochs, and trial condition normalization, 38.8% of trials were excluded. The data from all participants were analyzed.

### 3.3. Factorial Analysis

Our research question consists of two parts: What is the timing of structure sensitivity, relative to the timing of sensitivity to lexical content or agreement? What role does lexical processing play with respect to the SSE? For these reasons, we conduct two sets of analyses. The first analysis contrasts the Grammatical, Consonant String, Scrambled, and Ungrammatical conditions. The second pair of analyses then only includes the Grammatical trials to determine when we observe correlations between EEG signal and lexical variables.

Both sets of analyses use a two-stage regression analysis. At the first level, within each subject, a linear model is fit to each time point and sensor. For the factorial analysis, this model includes an intercept, and three β coefficients corresponding to the difference between Grammatical trials and Consonant String, Scrambled and Ungrammatical trials for each sensor and time point.

At the second stage, we conduct an across-participants analysis in which, for each effect separately, the β values are entered into a two-tailed one-sample *t*-test to determine, at each time point and each sensor, whether their values are significantly different from a population mean of 0 across participants. This produces *t*-values at each time point and sensor,. For estimating significance at the group level, we perform spatio-temporal cluster-based permutation testing over the resulting *t*-values (Maris & Oostenveld, 2007). This is a non-parametric method for first identifying clusters of significant responses that are contiguous in space and time, and then bootstrapping a null distribution from the data to estimate the statistics of the cluster. This allows for correcting for multiple comparisons, while acknowledging the non-independence of adjacent sensors or time points in an EEG recording. Adjacent time points and sensors (of the same polarity) were clustered together if their (uncorrected) *p*-value was *p* < 0.01. Cluster-forming *p* is a parameter which controls cluster size in space and time, rather than determining statistical significance. *T*-values were summed to calculate a cluster-level statistic. A null cluster distribution was estimated by repeating this procedure for 10,000 permutations in which the β value of each participant was randomly shuffled with 0. Cluster *p*-value was equated with position in the null distribution with the bottom 2.5% and top 2.5% being considered statistically significant. For the clustering procedure, we used a stricter cluster forming *p*-value threshold than may be typical (i.e., *p* < 0.05), because lower *p*-values favor smaller clusters, which may be more useful for estimating the temporal onset of effects. Additional *post-hoc* analyses with different cluster forming *p*-value thresholds (*p* < 0.05, *p <* 0.10) did not significantly change the pattern of results, and are thus not reported here.

Strictly speaking, spatio-temporal cluster-based permutation tests are only null hypothesis significance testing procedures for the size of a cluster in space-time, not for the temporal and spatial dimensions of the cluster (Rousselet, 2019; Sassenhagen & Deschkow, 2019). For this reason, we report the largest cluster (i.e., the smallest *p*-value). Afterwards, for each factorial comparison, we average the raw signal from the spatio-temporal coordinates of the cluster and conduct pairwise *t*-tests between Grammatical, Scrambled, Ungrammatical, and Consonant String. We also calculate and plot the correlation of the unigram and contextual probability of content words (words 2, 3, and 5; see § 3.4) in the Grammatical trials with the single-trial EEG data. This analysis also follows Fallon & Pylkkänen’s (2024) and Flower & Pylkkänen’s (2024) strategy of using the SSE as a spatio-temporal localizer followed by secondary analyses on what other effects are observable in the SSE cluster.

For the Consonant String comparison (Grammatical as reference), the largest cluster was positive-going and consisted of 32 centro-parietal sensors, 167–800ms, *p* < 0.001 (Fig. 2A). This cluster suggests a relatively uniform sustained difference in evoked response between Grammatical and Consonant String stimuli; the latter are immediately detectable as non-lexical, and thus none of the subsequent linguistic processes are likely to be engaged, unlike the other three conditions. The onset of this cluster falls within the N170+N250 complex, and the centro-parietal distribution is consistent with the topography of this ERP. Post-hoc analyses in this time window showed more positive activity for Consonant Strings compared to Grammatical, Scrambled, and Ungrammatical trials. There was a correlation of word 3 (main verb) contextual probability within the grammatical trials during this cluster (Pearson’s *r* = 0.29, *p* = 0.05), but no significant correlations between the contextual probability of the other lexical items nor unigram probability of any words.

**Figure 2.**
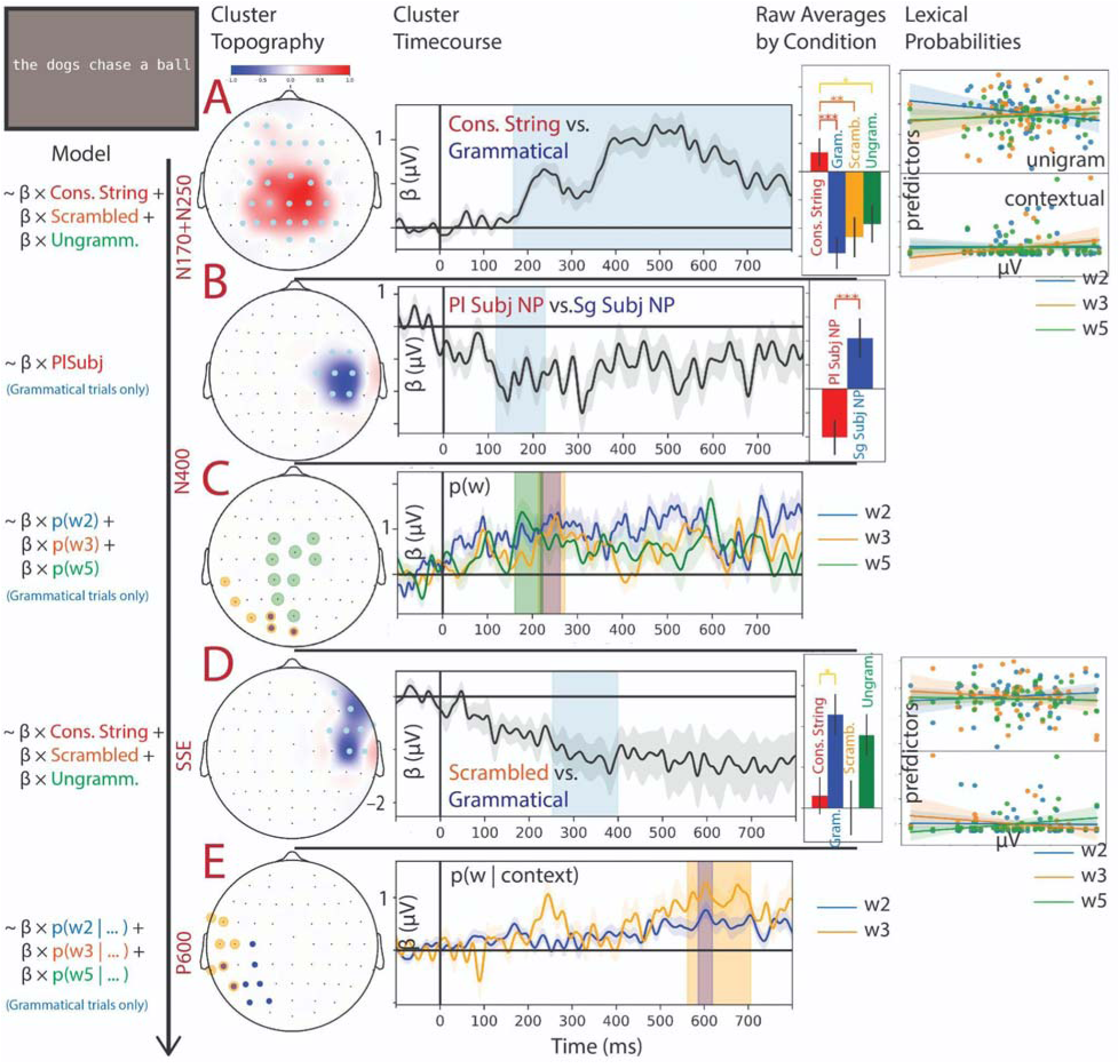
Clusters, pairwise activation comparisons, and correlations from the two-stage EEG analyses. (A, B, D) Topographic plots show the distribution of *t*-values and sensors included in each cluster. Time series show the average β coefficient value in these sensors over time, with the temporal extent of the cluster shaded. Bar plots show the average activity by condition in the spatial and temporal extent of the cluster (Grammatical vs. Scrambled vs. Consonant String vs. Ungrammatical for A and D; Singular vs. Plural Subject noun phrase (NP) in B). Error bars show one standard error above and below the mean; starred pairwise comparisons are significant at *p* < 0.05 with Hochberg correction for multiple comparisons. Scatterplots in A and D show the correlation of unigram probability (top) and contextual probability (bottom) for the Grammatical trials in the spatial and temporal extent of the cluster for word 2 (blue), word 3 (orange), and word 5 (green). (C, E) Topographic plots show the sensors included in each (overlapping) cluster for word 2 (blue), word 3 (orange), and word 5 (green) unigram probability (C) and contextual probability (E). Time series show the overlapping average β coefficient value in these clusters over time, with the temporal extent of the cluster shaded.

The Scrambled condition (Grammatical as reference) was associated with one significant cluster, which was negative-going and consisted of 8 right lateral sensors, 253–399ms, *p* = 0.036 (Fig. 2D). This cluster showed a sustained negative response for the Scrambled condition. The topography of this cluster is centered on anterior and right lateral sensors, although normally N400 effects are observed over centro-parietal and posterior sensors (cf. Wen et al., 2019). However, this time window is approximately the same as Wen et al.’s (2019) N400 finding. Pairwise comparisons between conditions revealed a significant difference between Grammatical and Consonant String trials (*p* < 0.05), but not between Grammatical and Ungrammatical trials. Within the grammatical trials, there were marginal correlations of word 3 (main verb) and word 5 (object noun) contextual probability during this cluster (Pearson’s *r* = 0.25, *p* = 0.08; Pearon’s *r* = 0.25, *p* = 0.09), but no significant correlations between the contextual probability of the word 2, nor the unigram probability of any words.

The Ungrammatical condition (Grammatical as reference) was not associated with any significant clusters (the largest having *p* = 0.331) and is not presented here. This was the case with clustering thresholds of *p* < 0.01, *p* < 0.05, *p* < 0.10, or *p* < 0.30. Thus, the failure to identify a significant cluster is not likely to be due to the stringentness of our clustering parameters.

To determine whether the lack of a difference between the Grammatical and Ungrammatical conditions indexes a failure to notice the singular vs. plural morphology (i.e., *-s,* vs. *-Ø* on the subject noun and the verb), we conducted a *post hoc* two-stage regression analysis with the same parameters comparing *singular* Grammatical trials (*the dog<u>Ø</u> chase<u>s</u> a ball*) and *plural* Grammatical trials (*the dog<u>s</u> chase<u>Ø</u> a ball*). Before a participant can realize that a stimulus is ungrammatical, they must first evaluate the number specification on both the subject noun phrase (NP) and verb, and then ‘check’ whether these values are appropriate given the syntactic structure of the sentence.

A significant cluster was observed consisting of 7 central right electrodes, 117–226ms, *p* = 0.03, (Fig 2B). Afterwards, we conducted *post hoc* Grammatical vs. Ungrammatical analyses nested within the singular trials and the plural trials. We did this because some behavioral findings demonstrate asymmetrical sensitivity to number errors in subject-verb agreement for plural controllers vs. singular controllers (Wagers et al., 2009). We failed to find any significant clusters distinguishing Grammatical vs. Ungrammatical, nested within singular or plural trials. Thus, while the EEG data indicate that participants distinguish trials with singular vs. plural subject NPs, neural responses are not sensitive to mismatch between subject NP and verb number specification.

### 3.4. Single-Trial Analyses

To determine the relative timing and topography of the SSE relative to activation of lexical items, we conducted two-stage linear regression analyses over unigram probability and contextual probability of the content words (*dogs*, *chase*, *ball*) in the Grammatical trials. This also permits us to determine whether individual lexical items are processed in parallel during parallel presentation of sentences. We conducted two separate single-trial analyses with two different measures of word likelihood: unigram frequency and BERT masked language modeling probability.

The analyses for the unigram and contextual probability analyses used the same two-stage regression procedure described in § 3.3. Both analyses included the continuous probability estimates for words 2 (subject noun), 3 (main verb), and 5 (object noun) in the same regression. The first and fourth words were not analyzed, (1) because they are contentless determiners (*the*, *a*); and (2) because, depending on which trials were excluded for a given participant, it was commonly the case that the only remaining trials available for analysis had a constant value for the first or fourth word position, i.e., all *the*’s in the first position or all *a*’s in the fourth position. Constant-valued regressors cannot be included in a linear model. Predictors were standardized before model estimation (i.e., shifted to mean zero and scaled to unit variance).

Unigram probability was calculated as the base 10 logarithm of the case insensitive frequency in the Google Books Ngram Dataset (v3, Michel et al., 2011). Contextual probability was estimated using BERT large, uncased, with whole word masking (Devlin et al., 2019), as accessed through Hugging Face (Wolf et al., 2020). The model had 336M parameters, with 24 layers, 16 attention heads, and with hidden dimensionality 1,024. Differing slightly from vanilla BERT, this BERT used whole word masking, in which rather than masking (potentially) sub-word units, all of the tokens corresponding to a word would be masked. We do not perform any comparisons, but we hypothesized that training to predict all of the sub-word tokens which make up a word would lead to more cognitively realistic whole-word prediction probabilities than the standard masked language modeling objective. Given an input with a mask (*the [MASK] chase a ball*), BERT produces a probability distribution over the tokens of its vocabulary. Thus, by masking words 2, 3, and 5 of each Grammatical stimulus item, we extract the model probability of each word in its bidirectional context.

For the unigram frequency analyses, we observed a significant cluster for the unigram probability of word 5 (the object noun), centered over 9 centro-parietal sensors, 161–223ms, *p* = 0.04. We also observed a marginally significant cluster for the unigram probability of word 3 (the main verb), centered over 6 left posterior sensors, 213–272ms, *p* = 0.08. These clusters are spatially adjacent, overlap temporally, and have a typical time and topographic location of an (early) N400 response. They are also positive-going, consistent with a reduction in the N400 for more probable words. We also report on the largest cluster observed for the unigram probability of word 2 (the subject noun), centered over 3 posterior sensors, 218–262ms, *p* = 0.17. This cluster shares the same wave morphology, topographic distribution, and temporal structure as the other unigram probability clusters. Taken together, this suggests that in rapid parallel reading, participants can coactivate multiple lexical items, each affecting the N400 independently. These three clusters are plotted together in Fig. 2C.

For the contextual probability analysis, we observed a cluster for word 3 (the main verb) centered over 7 left lateral sensors, 562–705ms, *p* < 0.01. We also observed a non-significant cluster for word 2 (the subject noun) centered over 8 left parietal and posterior sensors, 586– 618ms, *p* = 0.13. As before, these clusters both overlap in space and time, and both show an increase in positive activity for more contextually probable words, shown in Fig. 2E. These clusters overlap in time with typical ‘P600’ effects. We did not observe any clusters for word 5 (the object noun).

## 4. Discussion

Despite the overwhelming emphasis on serial, word-by-word reading in sentence processing research, fluent readers are capable of extracting abstract grammatical information from a single glance. How is this possible? Here, we contribute to the growing literature examining brain responses to grammatical properties of written sentences displayed rapidly in parallel (Asano & Yokosawa, 2011; Dufau et al., 2024; Fallon & Pylkkänen, 2024; Flower & Pylkkänen, 2024; Krogh & Pylkkänen, 2024; Massol et al., 2021; Snell & Grainger, 2017; Wen et al., 2019). We found that EEG responses in fluent English readers showed evidence of detecting linguistic vs. non-linguistic orthographic representations at ∼160ms post-sentence onset, and whether the arrangement of the words maps onto a syntactic representation by ∼300ms post-sentence onset. Moreover, we demonstrated that EEG signals correlated with unigram probability for content words ∼200–300ms, slightly earlier than the SSE. We also failed to find evidence that readers detected subject-verb agreement errors in English, either in the whole epoch or during the SSE space-time. Interestingly, we demonstrated distinct neural signatures for singular vs. plural subject NPs in the pre-SSE time period. Thus, although readers appear to ‘detect’ number morphology, they may not ‘check’ whether subject-verb agreement is grammatical in rapid parallel reading. Finally, we showed that the SSE likely precedes integration of the words into a larger syntactic/semantic representation, as benchmarked by the later, ‘P600’ effect of contextual lexical probability.

Taken together, we suggest that these findings support (our interpretation of) **the constituent structure hypothesis.** When reading a short sentence displayed rapidly, readers initially map the visual stimulus onto a letter-form or morpho-orthographic representation, during the M170/N170/N250 (Zweig & Pylkkänen, 2009; Grainger & Holcomb, 2009). In our study, this is reflected as the rapid distinction between the Consonant String stimuli and the Grammatical stimuli, and the rapid detection of subject NP number. Shortly thereafter, readers activate the lexical representations of the words. This is reflected in the N400(-style) effect reported in our unigram frequency analysis, with a reduction in negative activity from ∼200– 300ms for more probable words (Bornkessel-Schlesewksy & Schlesewsky, 2019; Kutas & Hilyard, 1984; Lau et al., 2008). Interestingly, we observed spatially and temporally overlapping clusters for each of the three content words in our target stimuli, which may suggest a parallel reading strategy. Following this, we observe the SSE, the detection of Grammatical trials vs.

Scrambled counterparts, ∼300ms. This ERP effect is temporally overlapping with the N400, but spatially distinct, centered over right anterior and lateral sensors. Within-cluster analyses failed to identify a difference between Grammatical trials and Ungrammatical trials, reflecting an insensitivity to subject-verb agreement processing (Fallon & Pylkkänen, 2024). Finally, we observe a pattern of ERP responses to the contextual probability of the content words, much later than the SSE, 500–800ms. Although the topographic distribution and wave morphology are not typical of a P600, the finding that contextual properties do not correlate with EEG responses until this late are consistent with models of the P600 as an index of integration difficulty (e.g., Kuperberg, 2007; Friederici, 2011). Crucially, the finding that the ERP corresponding to the SSE does not index either contextual integration nor subject-verb agreement are compatible with **the constituent structure hypothesis** (c.f., Fallon & Pylkkänen, 2024).

### 4.1. Lexical Access in RPVP

In our study, we observed one significant cluster corresponding to the unigram frequency of the main verb, a marginally significant cluster for the subject noun, and a non-significant cluster for the object noun. This pattern is suggestive of parallel reading of multiple words in a single fixation. Flower & Pylkkänen (2024) report temporally overlapping responses for *bigrams* in 4-word grammatical sentences (*all cats are nice*) at ∼250ms, contemporaneous with the early-stage SSE in their study. This may demonstrate an intermediate ‘chunking’ of words into units, which may be compatible with either coactivation of lexical and syntactic representations (Asano and Yokosawa, 2011; Snell and Grainger, 2017), or rapid identification of constituents.

Both our results and Flower & Pylkkänen’s (2024) results depend on a two-stage regression analysis. As discussed in § 3, the first stage of regression collapses across individual trials, and the second stage of regression collapses across participants at the group level. It is still feasible that, trial-to-trial, participants coactivate only one or two lexical items, if this is the maximum that is possible in a single (foveal) fixation (Brothers et al. 2017; White et al. 2019).

Additionally, we mention that our two-stage regression analysis only revealed one significant cluster post-correction. We therefore remain agnostic about the degree to which parallel reading is a component of comprehending sentences in ‘at-a-glance’ reading generally.

### 4.2. The SSE ERP

One concern about research on the SSE is that participants may be able to use their top-down knowledge of syntax while performing the task (match/mismatch or post-cued word recall) to make an informed guess for grammatical, but not ungrammatical trials, thus producing the behavioral sentence superiority effect (Staub, 2023). This account, however, is unable to explain observed EEG (e.g., here and in Wen et al., 2019) and MEG (e.g., Fallon & Pylkkänen, 2024; Flower & Pylkkänen, 2024; Krogh & Pylkkänen, 2024) SSEs in RPVP.

Another point is that the EEG SSE observed here could potentially be explained by selective fixation and differences in bigram frequency or transition probability between the conditions. Given the Grammatical stimulus *the dogs chase a ball* and the Scrambled stimulus *a dogs chase ball the*, if the participant were to fixate on a single word position for each trial e.g., the fourth, with the following word available parafoveally, they would process *a ball* for the Grammatical condition and *ball the* for the Scrambled condition. On average, the Grammatical items are likely to have 2-word substrings with higher bigram frequency or transition probability than the corresponding Scrambled items. The behavioral data, however, disprove a strong fixation account.

If participants used this strategy, then participant accuracy should be close to 40%, given that the position in which a ‘mismatch’ might occur is counterbalanced. Even if they focused on a single position and parafoveally processed the next word as well as the previous word, i.e., attended to three words in parallel without engaging in linguistic processing, then accuracy would be closer to 60%. Observed accuracy, however, is 90% for the Grammatical condition and 81% for the Scrambled condition. Further, if participants used this strategy, then we would predict that accuracy should be similar across conditions. But, we observe that accuracy is significantly higher for the Grammatical, Ungrammatical, and Scrambled conditions compared to the Consonant String conditions. Accuracy on the Consonant String conditions was approximately around this 60% benchmark, suggesting that participants may have used this kind of reading strategy in these non-linguistic cases. Lastly, this explanation seems unlikely given the lack of sensitivity to agreement errors, since it would be expected that the bigram and transition probabilities for agreement violations (*dogs chases*) would be lower than for grammatical relations (*dog chases*).

Secondly, we note some differences between the SSE ERP we elicited in our study vs. that of Wen et al (2019). Like their study, we observe an ERP for Grammatical trials compared to their Scrambled counterparts ∼250–400ms, in a typical N400 time window. Like them, we also observe a positive (= reduction in negative) response. However, our topographic distribution is over right anterior sensors, whereas theirs has a larger centro-parietal scalp distribution, more typical of the N400. The SSE response observed here is comparable to the MEG findings by Dufau et al. (2024), which show the neural SSE emerging in anterior portions of traditional language cortex at ∼320ms. By contrast, Fallon & Pylkkänen (2024) observe much earlier SSEs, around ∼125ms and ∼200–250ms. One consideration is that our stimuli had 5 words (3 content words, 2 functional words), Dufau et al.’s (2024) study used 4, and Fallon & Pylkkänen’s (2024) used 3 (all content words). It is possible that length may affect the latency of the SSE, especially if readers rely on a parallel reading mechanism; i.e., 3 short words may be more likely to be read foveally compared to 4 or 5 words. It could also suggest a degree of serialism such that the SSE is delayed as a function of how many words make up the syntactic tree.

### 4.3. Agreement and Contextual Integration in the SSE

The failure to detect agreement violations supports an interpretation in which the SSE reflects a rapid sketch of a syntactic representation rather than a detailed analysis. In ERP research, subject-verb agreement errors elicit a strong, early response (a LAN followed by a P600) in serial reading experiments, which suggests both detection of the ungrammaticality followed by a repair attempt (Angrilli et al., 2002; Coulson et al., 1998, Osterhout & Mobley, 1995; see Molinaro et al., 2011, for a review). Furthermore, if we accept that readers may process (some of) a short sentence in a single glance, why were they unable to notice the mismatch in number specification of two adjacent words in our study?

We suspect that there are two possible explanations for this. The first is that readers may deploy top-down grammatical knowledge to constrain or ‘correct’ the percept. Flower & Pylkkänen (2024) demonstrate that while the earliest phase of the neural SSE is sensitive to local transpositions ∼250–300ms (*all cats are nice*, *all are cats nice*), the later phase is not ∼300– 400ms. They interpret this finding as demonstrating an initial bottom-up detection of the error followed by top-down correction. The second possible explanation is that the mechanisms subserving the SSE do not permit integration of lexical items into a detailed analysis, in which relations between words are evaluated and interpreted. In either case, it is unclear why the later components of our ERP did not reveal sensitivity to the ungrammatical agreement relation, particularly during the 500–800ms stage in which we argue lexical items are integrated into the larger analysis.

Another possibility, however, is that comprehenders may simply ignore morphosyntactic information that is not conducive to identifying the semantic ’gist’ of a sentence (Asano & Yokosawa, 2011), which may be consistent with ’good enough’ parsing models (Ferreira & Patson, 2007). Subject-verb agreement in English is a formal grammatical dependency, which serves as an indirect cue to the interpretation of the sentence, which is arguably more important in ‘at-a-glance’ reading. Investigating whether the SSE is sensitive to selectional restrictions (*they will bring a book* vs. *they will bringing a book*; *they depend on him* vs. *they depend in him*) may further clarify why readers ignore agreement. We also observe that psycholinguistic research has long observed that agreement violations are particularly ‘fallible’, compared to other kinds of long-distance dependencies, such as anaphora (Wagers et al., 2009; Phillips et al., 2011; Dillon et al., 2013).

Lastly, the source of insensitivity to agreement may be due to the relatively weak visual salience of the number affixes in English (*-s*, -Ø). This could be further tested by contrasting more visually salient agreement errors, such as *child **is*** vs. *child-**ren are*** or leveraging agreement morphology in other languages with more complex and visually salient agreement morphemes (*asa-**vara*** ‘I love him’; *asa-**vassi*** ‘we love you,’ Kalaallisut/West Greenlandic; ILDD *kôr-**i*** ‘I do’; ILDDD *kôr-en* ‘you do,’ Bangla/Bengali).

Importantly, we found that readers do exhibit distinct ERPs for singular vs. plural subject NPs in grammatical trials, which suggests that comprehenders do not always fail to notice small suffixes like *-s*. This is to be expected if readers attend to morphological cues (‘interpretable features’) that are semantically relevant, such as number on subject NPs, but ignore morphological features (‘uninterpretable features’) that are semantically irrelevant. Investigating whether the neural SSE is sensitive to plurality in contexts that are semantically inert (i.e., *pluralia tantum* nouns like *scissors*), or agreement phenomena in languages that may be more semantically informative (i.e., number features in a null subject language) may be instructive.

Our study suggested distinct phases of lexical processing in the RPVP task – unigram probability corresponding to neural activity ∼200–300ms, and contextual probability ∼500– 800ms. As before, we acknowledge that, of the reported clusters, two are significant and one is only marginal.

While myriad work has related word-by-word information-theoretical complexity metrics from causal (left-to-right) language models (see Hale, 2016 for a review) to observed human data (e.g., self-paced reading time, EEG, fMRI) during language processing, we believe the use of probability from a *bidirectional* language model to be a novel application made possible by the use of RPVP. Future work remains to further investigate how predictors derived from bidirectional models can be related to language processing.

## 5. Conclusion

Our theories of language comprehension must be as flexible as the range of forms that language can take. In the visual domain, language can be understood slowly or from a quick glance – when editing a journal article vs. checking a text notification. Here, we demonstrate that there are several distinct stages involved during the processing of sentence-length written stimuli viewed only momentarily. After the rapid detection of linguistic content ∼160ms and initial activation of lexical items at ∼200ms, a ‘sketch’ of a grammatical analysis occurs at ∼300ms.

These findings are compatible with the hypothesis that the initial sketch involves identification of basic constituent structure (cf., Fallon & Pylkkänen, 2024). This initial sketch is insensitive to formal grammatical errors, such as subject-verb agreement errors, and precedes deeper interpretative processes, occurring after ∼500ms. We also demonstrate that several language-related ERPs are observed in the processing of sentences ‘at-a-glance’, and that coactivation of lexical items may be observable in a typical ‘N400’ response, just prior to the SSE.

## Notes

### Competing Interest Statement

The authors have declared no competing interest.

### Summary of Updates

methodological revisions with new result interpretations

